# A framework for predicting potential host ranges of pathogenic viruses based on receptor ortholog analysis

**DOI:** 10.1101/2020.12.07.414292

**Authors:** Guifang Du, Yang Ding, Hao Li, Xuejun Wang, Junting Wang, Yu Sun, Huan Tao, Xin Huang, Kang Xu, Hao Hong, Shuai Jiang, Shengqi Wang, Hebing Chen, Xiaochen Bo

**Affiliations:** Beijing Institute of Radiation Medicine, Beijing 100850, China

**Keywords:** Zoonotic Virus, Receptor Orthologs, Potential Host Prediction, SARS-CoV-2

## Abstract

Viral zoonoses are a serious threat to public health and global security, as reflected by the current scenario of the growing number of severe acute respiratory syndrome coronavirus 2 (SARS-CoV-2) cases. However, as pathogenic viruses are highly diverse, identification of their host ranges remains a major challenge. Here, we present a combined computational and experimental framework, called REceptor ortholog-based POtential virus hoST prediction (REPOST), for the prediction of potential virus hosts. REPOST first selects orthologs from a diverse species by identity and phylogenetic analyses. Secondly, these orthologs is classified preliminarily as permissive or non-permissive type by infection experiments. Then, key residues are identified by comparing permissive and non-permissive orthologs. Finally, potential virus hosts are predicted by a key residue–specific weighted module. We performed REPOST on SARS-CoV-2 by studying angiotensin-converting enzyme 2 orthologs from 287 vertebrate animals. REPOST efficiently narrowed the range of potential virus host species (with 95.74% accuracy).

## Introduction

Viral zoonoses pose serious threats to public health and global security, and have caused the majority of recent human pandemics [e.g., those of HIV, Ebola, severe acute respiratory syndrome (SARS), and avian influenza] (Kreuder Johnson et al., 2015; Olival et al., 2017; Stephen S et al., 2012). An understanding of the species tropism of viral transmission is thus key for the development of pandemic control programs. The global coronavirus disease 2019 (COVID-19) pandemic caused by severe acute respiratory syndrome coronavirus 2 (SARS-CoV-2, also, 2019-nCoV and COVID-19 virus) has caused an unprecedented public health and economic crisis (Zhou et al., 2020). Comparison with related coronavirus sequences has shown that SARS-CoV-2 likely originated in bats, followed by transmission to an intermediate host (Dos Santos Bezerra et al., 2020; Lam et al., 2020; Lopes, de Mattos Cardillo, & Paiva, 2020; Zhou et al., 2020). Numerous studies suggest that a diversified host range is involved in the SARS-CoV-2 pandemic outbreak (Abdel-Moneim & Abdelwhab, 2020; Hossain, Javed, Akter, & Saha, 2020). *In vivo* studies have demonstrated that mink, ferrets, cats, dogs, and some non-human primates are susceptible to SARS-CoV-2, whereas mice, white-tufted-ear marmosets, chickens, ducks, and tree shrews are not (Abdel-Moneim & Abdelwhab, 2020; Hossain et al., 2020; S. Lu et al., 2020; Oreshkova et al., 2020; Shi et al., 2020; X. Zhao et al., 2020; Y. Zhao et al., 2020). The SARS-CoV-2 outbreak is another critical example proving the existence of close, straightforward interaction among humans, animals, and environmental health that can result in the emergence of a deadly pandemic.

A large number of viruses, including coronavirus and influenza virus, has existed in nature for a long time (Cui, Li, & Shi, 2019; D. Liu, Ma, Jiang, & He, 2019; Olival et al., 2017; Tiwari et al., 2020; Xiaoman, Xiang, & Jie, 2020). These viruses live in and coexist with animals. They are also constantly mutating and awaiting opportunities to invade human beings. Animal surveys resulted in the discovery of many thousands of new viruses. Such research would benefit studies of viral diversity and evolution, and could determine whether and why some pathogens cross species boundaries more frequently than others (Bae & Son, 2011; Edward C, Andrew, & Kristian G, 2018; Lasso et al., 2019; Olival et al., 2017; Stephen S et al., 2012). Given the rarity of outbreaks, however, it is arrogant to imagine that we could use such surveys to predict and mitigate the emergence of disease. In addition, due to adaptive genetic recombination, the possibility that a new coronavirus or influenza virus will evolve cannot be excluded, as reflected by the current scenario of the growing number of SARS-CoV-2 cases. There is a small, but real, possibility that SARS-CoV-2 will take refuge in a new animal host and be reintroduced to humans in the future. Thus, the possibility of interspecies transmission of viral infections in hot spots is of concern to human beings. As these viruses are very diverse, evaluation of the threat that they pose remains a major challenge, and efficient approaches to the rapid prediction of potential animal reservoirs are needed.

One way in which virologists can attempt to predict potential host species is via the cellular receptors of viruses. The recognition of and interaction with cellular receptors are critical initial steps in the infectious viral life cycle and play key regulatory roles in host range, tissue tropism, and viral pathogenesis (Maginnis, 2018). In addition, the gain of function of a virus to bind to receptor counterparts in other species is prerequisite for interspecies transmission (G. Lu, Wang, & Gao, 2015). Human angiotensin-converting enzyme 2 (ACE2) has been identified as the cellular receptor for SARS-CoV-2(Hamming et al., 2004; Zhou et al., 2020). SARS-CoV (Wenhui Li et al., 2003) and human coronavirus NL63 (Hofmann et al., 2005; Kailang Wua, Weikai Lib, Guiqing Penga, & Lia, 2009) have caused human disease previously and interact with ACE2 to gain entry into cells. ACE2 is expressed in a diverse range of species throughout the subphylum Vertebrata. The host range of SARS-CoV-2 may be extremely broad due to the conservation of ACE2 in mammals (G. Lu et al., 2015). Using *in vitro* functional assays, Liu et al. (Y. Liu et al., 2020) showed that 44 mammalian ACE2 orthologs, including those of domestic animals, pets, livestock, and animals commonly found in zoos and aquaria – but not orthologs in New World monkeys – could bind the SARS-CoV-2 spike protein and support viral entry. In addition to performing receptor sequence analysis, Kerr et al. (Kerr et al., 2015) developed a combined computational and experimental approach to assess the compatibility of New World arenaviruses with potential new host species, although this method is suitable only for the rodent host range. Using the random forest machine-learning algorithm, Eng et al. (Eng, Tong, & Tan, 2014) constructed computational models for prediction of the host tropism of influenza A virus, but this method is suitable only for the avian and human host ranges. Many other computational approaches have been developed to predict bacteriophage–host relationships, but whether these methods can be applied to animal viruses is unknown (Ahlgren, Ren, Lu, Fuhrman, & Sun, 2017; Edwards, McNair, Faust, Raes, & Dutilh, 2016; Roux, Hallam, Woyke, & Sullivan, 2015).

With the advancement of high-throughput next-generation sequencing, the virus isolation technology that enables identification of the pathogen causing an outbreak is no longer the bottleneck it once was (Jonsdottir & Dijkman, 2016; Ko, Salem, Chang, & Chao, 2020; Zhu et al., 2020). Novel and traditional techniques have proven to be extremely useful for the discovery of viral receptors; in particular, biochemical and structural analyses have provided a great deal of insight into the molecular interactions between viruses and receptors (Maginnis, 2018). In addition, the continuous enrichment of protein sequence databases has increased the number of species for which receptor protein sequences are available. These excellent conditions enable the development of approaches for the prediction of potential hosts of pathogenic viruses. In this report, we describe a combined computational and experimental approach called REceptor ortholog-based POtential virus hoST prediction (REPOST), which provides a flexible framework for the identification of potential virus hosts based on virus receptor ortholog analysis. REPOST takes virus cellular receptor orthologs from multiple species as input and predicts the possibility of undetermined species’ roles as potential hosts. Using this framework, we first systematically analyzed ACE2 orthologs from 287 vertebrates, primarily mammals. Then, we analyzed the binding ability of ACE2 to viruses from 16 representative species in a pseudovirus infection experiment. Thereafter, we identified 95 key ACE2 residues that may destroy ACE2–SARS-CoV-2 interaction. Finally, we used a residue-weighted calculation method to predict the possibility of ACE2 as a receptor in unknown species. The accuracy of the framework is very high, as proven by the second functional experiment.

## Results

### Design and comprehensive features of the REPOST workflow

REPOST application requires the following dependences: (1) known cellular receptor of the virus, the availability of receptor orthologs from most species [we recommend downloading receptor protein sequences from the National Center for Biotechnology Information (NCBI) database], and ability to perform an *in vitro* virus or pseudovirus infection experiment (See Methods). The REPOST workflow is as follows (Fig. 1). (1) Receptor orthologs are selected for the following infection experiment based on identity and phylogenetic analyses among various species. The species of the selected orthologs should be representative of taxonomic groups (e.g., primates, rodents, carnivores, bats, marsupials, birds) and should better to be frequently in close contact with humans. And the orthologs should have high degrees of relative identity with those from known virus hosts. (2) Each selected ortholog is preliminarily classified as permissive or non-permissive (meaning that cells with the ortholog overexpression is or is not permissive of virus entry) by performing virus or pseudovirus infection experiment. In addition, published experimental evidence can also be used to supplement our classification. At least three orthologs of each type are needed. (3) Key residues are identified by comparing permissive and non-permissive orthologs (for details, see the SARS-CoV-2 example). (4) Potential virus hosts are predicted by a key residue–specific weighted module.

**Figure 1.**
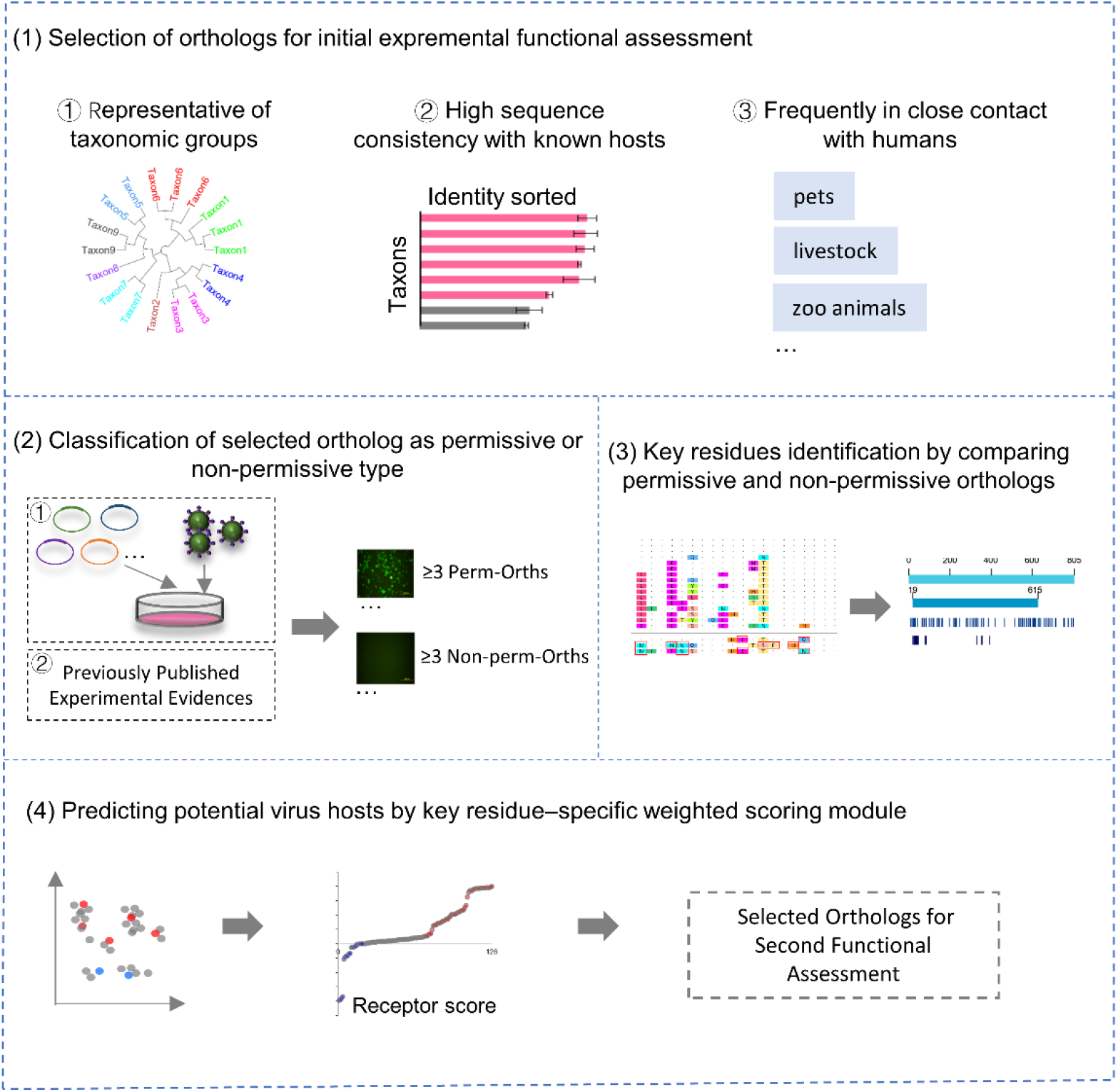
REPOST design and workflow.

Taken together, based on a combination of virus infection experiments and computational analysis, REPOST can be used to extrapolate, on a large scale, the possibility of undetected species being virus hosts.

### Application of REPOST to predicting potential host range of SARS-Cov-2

#### Selection of ACE2 orthologs from various species by identity and phylogenetic analysis

We collected the DNA, mRNA, and protein sequences of ACE2 orthologs from 287 vertebrate animals from the NCBI database. These species were mammals (n = 126), birds (n = 73), amphibians (n = 4), bony fishes (n = 66), lizards (n = 9), and other chordates (n = 10). The mammals were primates (n = 26), rodents (n = 23), carnivores (n = 23), whales and dolphins (n = 10), bats (n = 13), even-toed ungulates (n = 14), marsupials (n = 4), and other placental species (n = 13) (Fig. 2A). The length ranged from 6694 to 107639 bp for DNA sequences, and 1035 to 7004 bp for mRNA sequences (Fig. S1A, B). Protein sequences contained 344 to 862 amino-acid residues, and the length in 25% of species was the same as that of human (805 amino-acid residues) (Fig. S1C).

**Figure 2.**
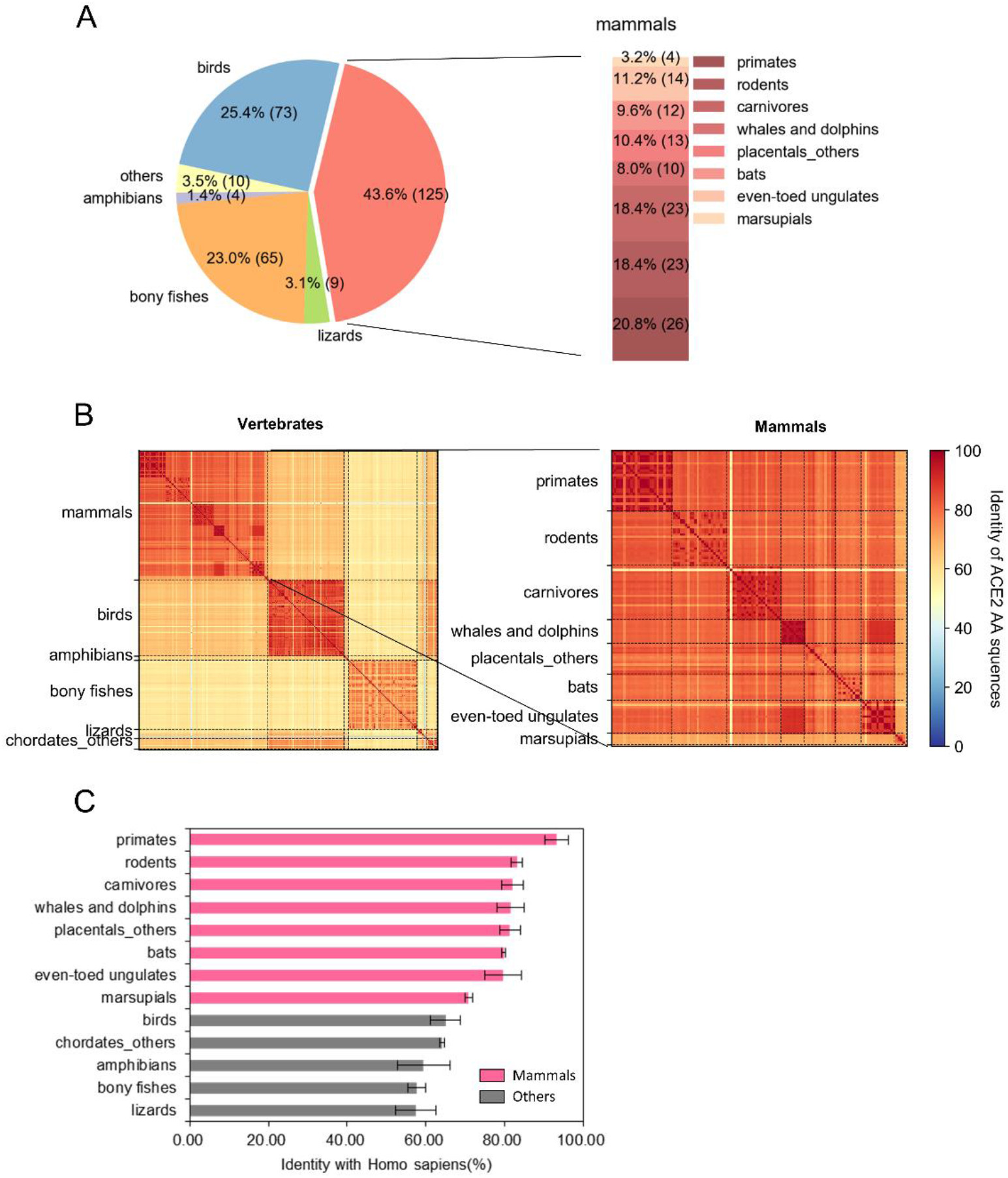
Identity analyses of ACE2 orthologs from 287 vertebrates. (A) Amino-acid sequences of ACE2 in 287 vertebrates from the NCBI dataset. Bar of pie chart showing the numbers and proportions of species in each group. (B) Left: ACE2 identity matrices for 126 mammals. Right: enlargement of a portion of the left panel. (C) Ranked consistency of ACE2 among all species to humans. Values are expressed as means of identity for each taxon with standard deviations (SDs; error bars).

We first performed the phylogenetic analysis, and found that ACE2 orthologs from the same taxon were usually clustered into the same branch. (Fig. S2). Also, we analyzed the identities of all sequences pairwisely, the result indicate that the ACE2 protein sequences were highly conserved across each taxon examined, as well as each subclass of mammals (Fig. 2B), suggesting that we can start classification with a few representatives from each taxon. Then, we ranked the identity of ACE2 among all species taxon to humans, and the result yielded the following order from high to low was as below: primates, rodents, carnivores, other placental species, even-toed ungulates, whales and dolphins, bats, marsupials, birds, other chordates, amphibians, lizards, and bony fishes (Fig. 2C). Based on the ranking result, we found that ACE2 in other vertebrate species except mammals had low consistency with that of human, suggesting they were not likely to be potential hosts or reservoirs for SARS-CoV-2. Previous study has supported our observation that poultry (belong to birds) is not susceptible to SARS-CoV-2(Shi et al., 2020). We finally chose 16 representative ACE2 orthologs (very similar to that of humans) from mammals (include primates, rodents, carnivores, bats, and even-toed ungulates) for following analysis. (Fig. 3). Among these species are wild animals, zoo animals, pets, and livestock that are frequently in close contact with humans, and model animals used in biomedical research.

**Figure 3.**
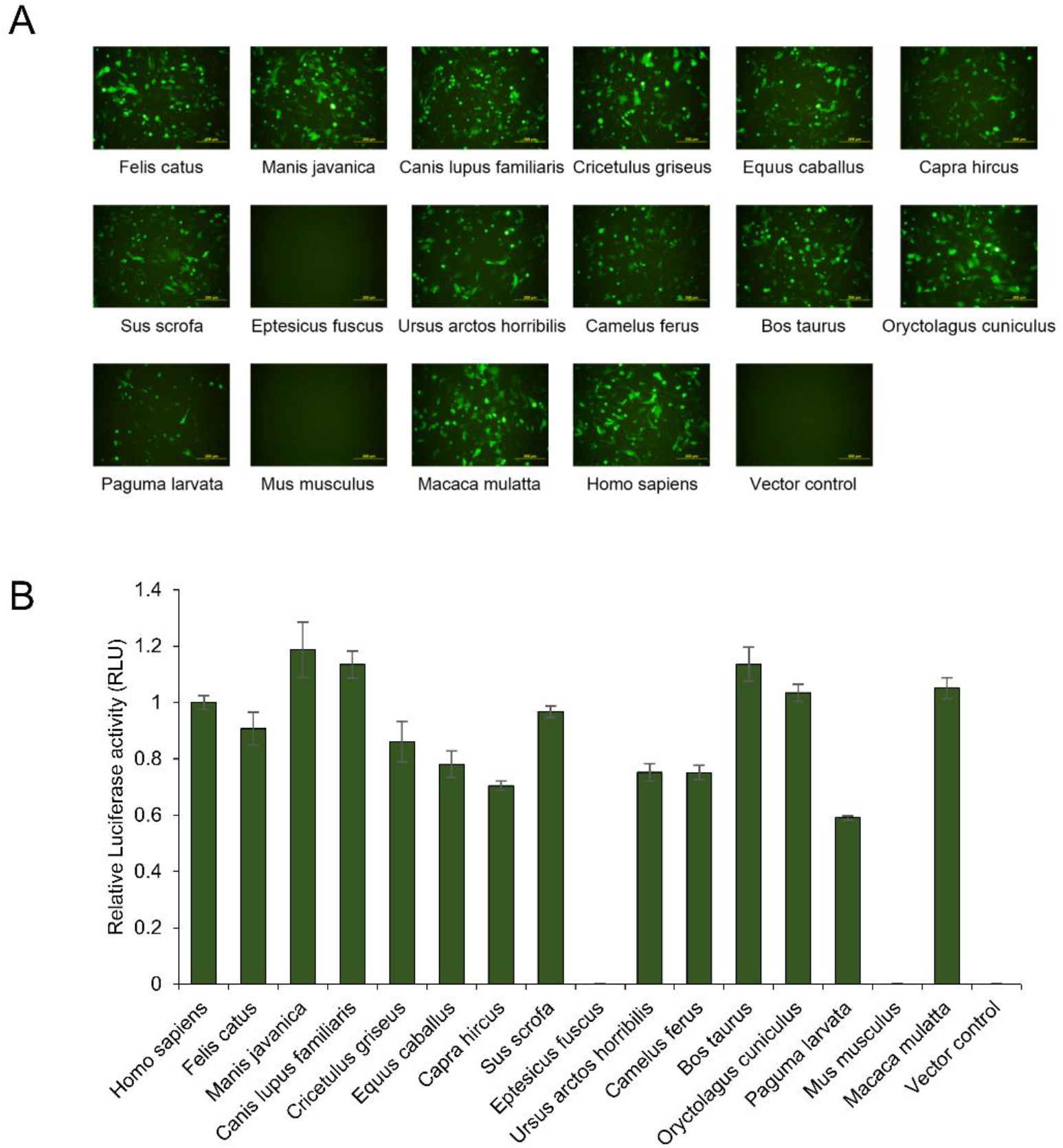
Functional assessment of ACE2 orthologs mediating SARS-CoV-2 pseudovirus particle (SARS2pp) entry. BHK-21 cells transfected with ACE2 orthologs or empty vectors were infected with SARS2pps (MOI = 3). (A) Expression of the eGFP protein, visualized by fluorescence microscopy. (B) Luciferase activity of different ACE2 orthologs in vector-transfected cells at 60 h after SARS2pp infection. Values are expressed as means with standard deviations (SDs; error bars).

#### Classification of permissive and non-permissive ACE2 orthologs by infection experiment in vitro

We infected BHK-21 cells ectopically expressing individual selected ACE2 orthologs with SARS-CoV-2 pseudovirus particles. As expected, BHK-21 cells lacking endogenous ACE2 expression were not permissive of SARS2pp infection. BHK-21 cells expressing ACE2s from *Homo sapiens, Felis catus, Manis javanica, Canis lupus familiaris, Equus caballus, Capra hircus, Sus scrofa, Ursus arctos horribilis, Camelus ferus, Bos taurus, Oryctolagus cuniculus, Paguma larvata* and *Macaca mulatta* were permissive of SARS2pp entry. ACE2s from *Eptesicus fuscus and Mus musculus* were not permissive of SARS2pp entry cells (Fig. 3A). The luciferase activity of different ACE2 orthologs of vector-transfected cells was consistent with the eGFP signals (Fig. 3B). Besides, we regarded ACE2 of *Rhinolophus sinicus, Chlorocebus sabaeus,* and *Macaca fascicularis* as permissive, and regarded *Callithrix jacchus* as non-permissive orthologs respectively, as published experimental evidence proved that ACE2 of *R. sinicus* could mediate SARS-CoV-2 entry into HeLa cells(Zhou et al., 2020), *Chlorocebus sabaeus(Cross et al., 2020),* and *Macaca fascicularis(S. Lu et al., 2020) were* susceptible to SARS-CoV-2 infection, and yet *C. jacchus* was not susceptible to SARS-CoV-2 infection(S. Lu et al., 2020).

Taken together, we eventually get 17 permissive and 3 non-permissive ACE2 orthologs. These orthologs would be used for the identification of key residues which changes of them may damage the ACE2-SARS-CoV-2 interaction.

#### Identification of key residues by comparing permissive and non-permissive ACE2 orthologs

After sequence comparison, we found 33 ACE2 protein residues of *M. musculus*, 20 residues of *C. jacchus*, and 50 residues of *E. fuscus* that differed from those of all SARS-CoV-2–permissive species (Fig. 4A, Table S1). Take the residues at the ACE2–SARS-CoV-2 interaction interface as an example, we found that substitutions in residues Q24, D30, K31, E42, M82, Y83, K353, and G354 that distinguished ACE2s of *C. jacchus*, *M. musculus*, and *E. fuscus* from those of all permissive species (Fig. 4B). These residues may be the key sites affecting the ACE2–SARS-CoV-2 binding.

**Figure 4.**
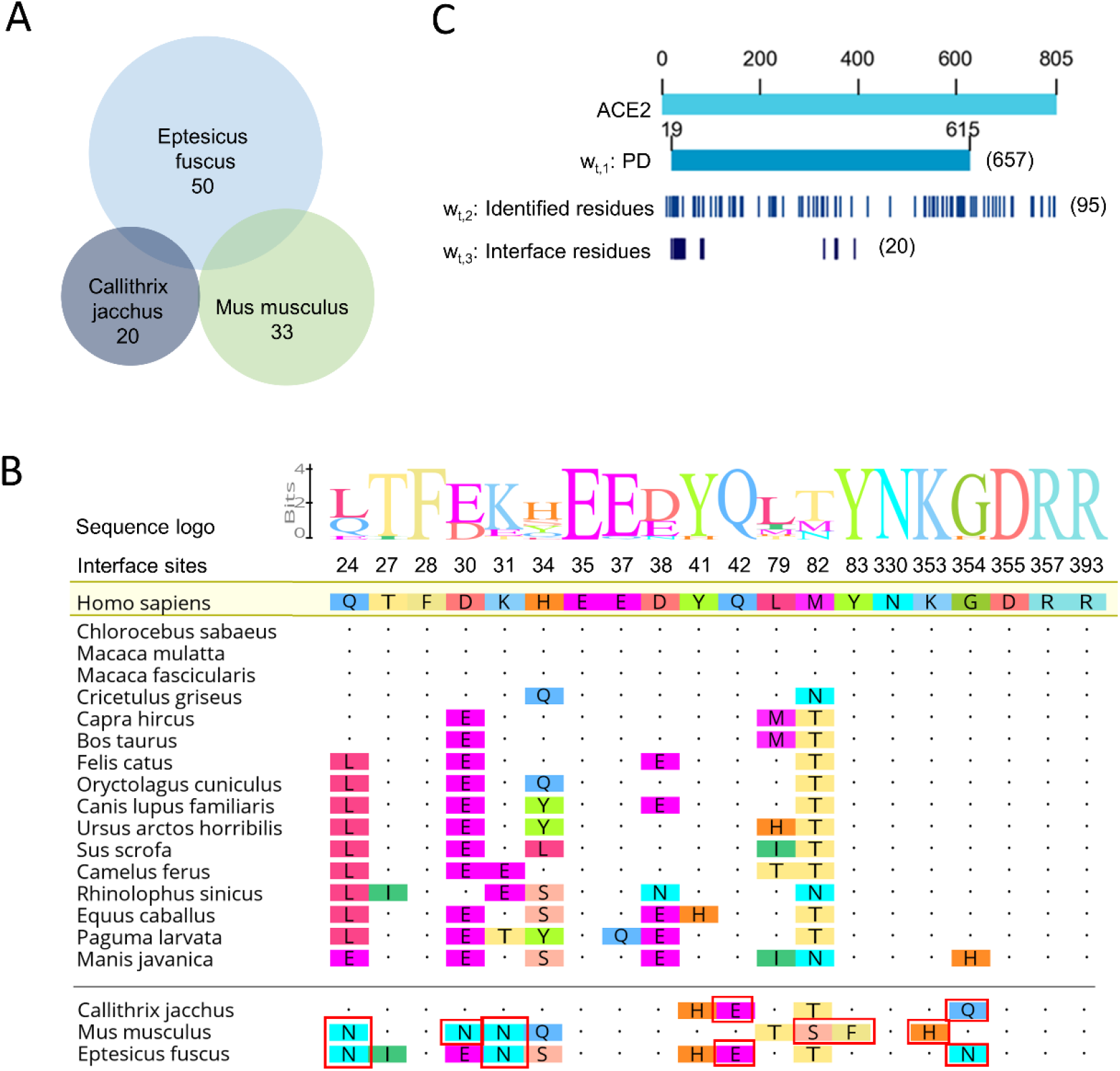
Identification of key residues and ACE2 residues at the interface with the viral spike protein. (A) Schematic depicting the identification of key differential ACE2 amino-acid sites, via the identification of distinctions between non-permissive ACE2 ortholog and all permissive orthologs. (B) Top: Sequence logos of the residues at S protein interfaces for all 17 permissive ACE2 orthologs. Bottom: Alignment of contacting residues from 17 permissive and 3 non-permissive ACE2 orthologs. Only amino acids that differ from those of humans are shown. Red boxes indicate ACE2 residues that may be key to the destruction of interaction with SARS-CoV-2. (C) Schematic diagram of the residues in the PD, the 95 key residues identified above, and the contact surface with the SARS-CoV-2 S protein of ACE2.

The ACE2 orthologs were characterized by a peptidase M2 domain (PD) which located outside the cell membrane and mediates virus binding, and a collectrin domain. In PD, there are 20 amino-acid residues located in the ACE2–SARS-CoV-2 contact interface (Lan et al., 2020; Renhong Yan et al., 2020; Wang et al., 2020; Wrapp. et al., 2020), and these residues are critical for virus recognition (Lan et al., 2020). The 75 of identified residues were located in the PD and 8 were located at the ACE2–SARS-CoV-2 interface (Fig. 4 B, C). We believed that the changes in amino acids at these sites might have a greater influence on the destruction of ACE2–SARS-CoV-2 binding than do changes at other sites.

#### Prediction of SARS-CoV-2 potential hosts by key residue–specific weighted module

Based on the above work, we developed a residue-specific weighted module for the prediction of susceptibility of untested mammal species (Figure 5A). The module takes as input multiple receptor orthologs, which including permissive, non-permissive orthologs from tested species and orthologs from other species to predict. It will first calculate a residue-weighted distance matrix for all orthologs taking into account three priors of the PD domain, the 95 key residues and the contact surface with SARS-CoV-2 S protein (Fig. 4C). We then used this distance matrix to select as potentially permissive (or non-permissive) candidates orthologs that were much closer to known permissive (or non-permissive) orthologs than to known non-permissive (or permissive) orthologs (Methods).

**Figure 5.**
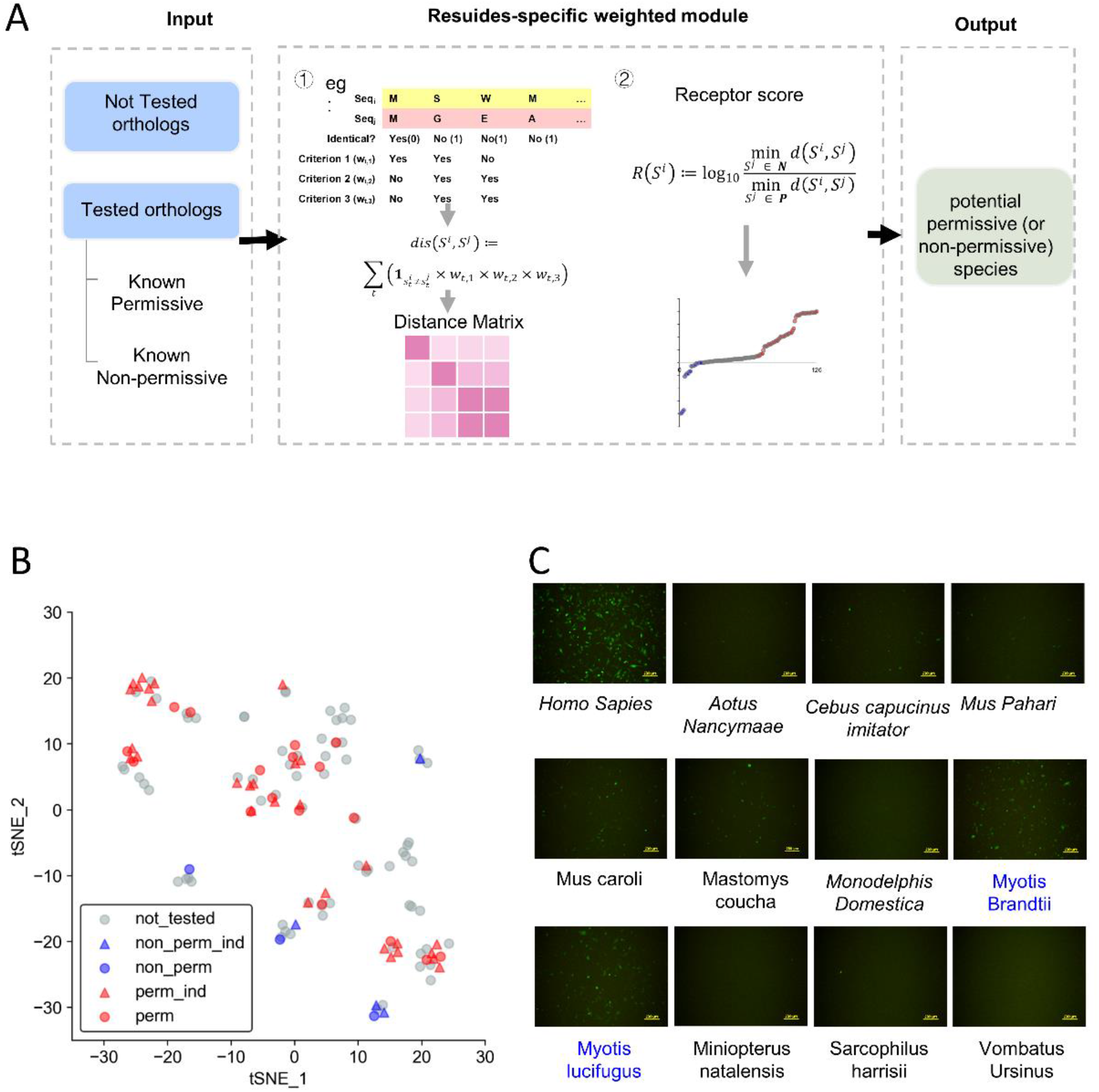
The residue-specific weighted prediction module. (A) Schematic diagram showing the workflow of the residue-specific weighted calculation model. (B) t-SNE projection plot showing the clustering results for ACE2 orthologs from 126 mammals. perm, ACE2 orthologs permissive of SARS2pp entry (see Figure 3); non_perm, ACE2 orthologs not permissive of SARS2pp entry (Figure 3); perm_ind, ACE2 orthologs permissive of SARS2pp entry according to independent research (Y. Liu et al., 2020); non_perm_ind, ACE2 orthologs not permissive of SARS2pp entry according to independent research (Y. Liu et al., 2020); not_tested, ACE2 orthologs that had never been tested. (C) Functional assessment of SARS2pp entry mediation of predicted non-permissive ACE2 orthologs.

The optimized distance matrix clearly separated known permissive and non-permissive orthologs, with no discernible mixing (Fig. 5B), as supported by independent experimental evidence (Y. Liu et al., 2020) (Fig. 5B; see also Table S2 for a full list of predictions for all 56 or 20 orthologs close to known permissive or non-permissive orthologs). Also, it displayed excellent performance in the extrapolation of non-permissive orthologs (which are currently scarce and potentially critical to the mechanistic understanding of SARS-CoV-2 tropism), with 9 of 11 identifications validated by subsequent assay of pseudovirus entry into BHK-21 cells (Fig. 5C). The other two orthologs showed only weak infection positivity (infection rates < 35%).

Taken together, REPOST, combined of phylogenetic analysis, experimental functional assessment, and residues-specific weighted module, provide an efficient framework for screening key residues of virus receptor that determine virus–receptor interaction and assessing the potential host ranges of pathogenic viruses. Compared with other methods based on molecular docking or simple molecular consistency analysis (Kerr et al., 2015; Y. Liu et al., 2020), REPOST has higher accuracy and a wider range of applications. In SARS-CoV-2 case, we found that ACE2 orthologs from a wide range of mammals, including pets, livestock, and animals commonly kept in zoos or aquaria, could act as functional receptors to mediate SARS-CoV-2 infection. Many of these species have been proved in previous experimental evidence (Abdel-Moneim & Abdelwhab, 2020; Hossain et al., 2020).This suggest that SARS-CoV-2 might have a broad host tropism and underscore the necessity to monitor susceptible hosts to prevent future outbreaks. In addition, we identified 10 previously unrecognized non-permissive ortholog sequences from primate, rodent, bat, and marsupial species. We also predicted that species other than mammals were not likely to be the host of SARS-CoV-2, as all non-mammalian ACE2 orthologs were too dissimilar from known permissive orthologs (Fig. S3A).

## Discussion

This study introduces and demonstrates the use of REPOST for the identification of potential virus hosts, using ACE2 receptor orthologs of SARS-CoV-2 as an example. This method showed a high host prediction accuracy. Our results suggest that SARS-CoV-2 cellular receptor ACE2 orthologs are strongly conserved across mammalian species, indicating the importance of the physiological function of ACE2. Furthermore, the protein sequence diversity of ACE2 was greater among bats than among other tested mammals, and ACE2 orthologs of bats were located on two distant branches of the evolutionary tree, highlighting the possibility that bat species act as reservoirs of SARS-CoV-2 or its progenitor viruses. Notably, we also found that ACE2 orthologs from a wide range of mammals, including pets (e.g., cats and dogs), livestock (e.g., pigs, cattle, rabbits, sheep, horses, and goats), and animals commonly kept in zoos or aquaria, could act as functional receptors to mediate SARS-CoV-2 infection when ectopically expressed in BHK-21 cells, suggesting that SARS-CoV-2 has a diverse range of hosts and intermediate hosts. These findings highlight the importance of the surveillance of animals in close contact with humans as potential zoonotic reservoirs. These results suggest that the potential host of the virus is related to the phylogeny of the species to a certain extent, but it is not accurate to predict based on this solely. Based on our experimental results, we identified 95 differences in amino-acid residues between SARS-CoV-2– permissive and –non-permissive ACE2 orthologs, eight of which are located at the ACE2–S protein interface. Some of these residues have been confirmed in other studies (F. Li, 2013; W. Li et al., 2005; Procko, 2020). The final prediction results were satisfactory and consistent with independent experimental evidence^28^, and the module showed excellent performance in the extrapolation of non-permissive orthologs. It is worth mentioning that the analysis identified 10 previously unrecognized non-permissive ortholog sequences from primate, rodent, bat, and marsupial species. These findings will enrich negative datasets, increasing the accuracy of the screening of key residues that affect virus–receptor interaction, and will aid the establishment and training of optimized predictive models.

We propose that REPOST will strengthen the ability to rapidly identify potential hosts of new pathogenic viruses affecting not only humans, but also animals. Another advantage of REPOST is the ease of key residue screening, which may lead to the identification of promising targets for the development of broad-spectrum antiviral therapies. REPOST can also be applied in other cases; for viruses with more than one cellular receptor, for example, all receptor orthologs can be used as input for systematic analysis. When the viral receptor cannot be identified, sequence information for all cellular membrane proteins can be integrated as input for prediction. REPOST, however, can be further improved. Other factors, such as transmembrane protease serine 2 (TMPRSS2) and the host immune response, can affect the susceptibility of animals (Bourgonje et al., 2020; Hoffmann et al., 2020). In some cases, viruses must open numerous “locks,” which can be conceptualized as a doorknob plus a deadbolt, to invade cells. In future research, we will explore the use of different sources of virus and host information to analyze the impacts of different characteristics on prediction results. In summary, the establishment of the REPOST predictive framework may be of great significance for the prevention and control of future outbreaks.

## Materials and methods

### Protein sequence identity and phylogenetic analyses

The amino-acid sequences of ACE2 orthologs from 287 vertebrates were downloaded from the National Center for Biotechnology Information (NCBI) database (https://www.ncbi.nlm.nih.gov/). Numbers in each sequence correspond to GenBank (https://www.ncbi.nlm.nih.gov/genbank/) accession numbers. ACE2 protein sequence identity, defined as the percentage of identical residues between two sequences, was analyzed using MEGA-X software (version 10.05) (Kumar, Stecher, Li, Knyaz, & Tamura, 2018) and the MUltiple Sequence Comparison by Log-Expectation (MUSCLE) algorithm (Edgar, 2004). Then, a phylogenetic tree was built using the minimum-evolution method with MEGA-X.

### Expression vector and cell lines

The cDNAs encoding ACE2 orthologs tagged with hexa-histidine were synthesized and cloned into a pCAG vector derived from pcDNA3 by replacement of the CMV promoter with the CAG promoter. All constructs were verified by Sanger sequencing. BHK-21 cells were maintained in Dulbecco’s modified Eagle medium (DMEM, Gibco) supplemented with 10% (vol/vol) fetal bovine serum (Gibco), 1 mM sodium pyruvate (Gibco), 1× non-essential amino acids (Gibco), and 50 IU/ml penicillin/streptomycin (Gibco) in a humidified 5% (vol/vol) CO_2_ incubator at 37°C.

### Pseudovirus infection of BHK21 cells expressing ACE2 orthologs

BHK-21 cells were seeded at 1 × 10^4^ cells per well in 96-well plates 12–18 h before transfection. The cells were then transfected with 100 ng control or ACE2 orthologs expressing plasmids with GenJet™ reagent (ver. II; SignaGen). The culture medium was refreshed 6–8 h after transfection. Another 6–16 h later, cells in each well were infected with SARS-CoV-2 pseudovirus bearing dual-reported genes (eGFP and luciferase) at an MOI of 3 or 10. The culture medium was changed 12 h after infection. At 60–72 h post-infection, images of eGFP expression were captured under a fluorescent microscope (IX73; Olympus) and luciferase activity was detected with the Steady-Lumi™ II Firefly luciferase assay kit (Beyotime).

### Details of the prioritization module

The weighted (Manhattan) distance for each pair of ortholog sequences *S*^*i*^, *S*^*j*^ was defined as

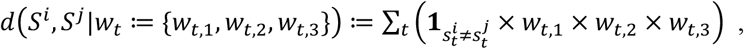

where 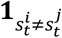 is the indicator function that takes 1 when the residues of *S*^*i*^ and *S*^*j*^ at the *t*th position of the alignment differ (i.e., 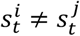) and 0 otherwise, and *w*_*t*_ ≔ {*w*_*t*,1_, *w*_*t*,2_, *w*_*t*,3_} are weights for the following three priors: (1) whether the residue at this position in the human ortholog is in the PD (*w*_*t*,1_)(Wang et al., 2020), (2) how much residue composition at this position differs between all permissive and non-permissive species from the experiment (*w*_*t*,2_), and (3) whether the residue at this position in the human ortholog is on the contact surface (*w*_*t*,3_)(Lan et al., 2020). For MUSCLE alignment, we used default parameters and removed all positions at which the human ortholog was gapped.

The optimal *w*_*t*_ was selected from the 1,000 possible combinations listed in Table S3 to maximize the following separation score:

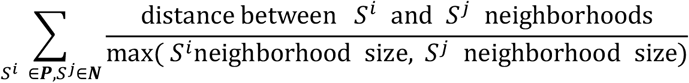

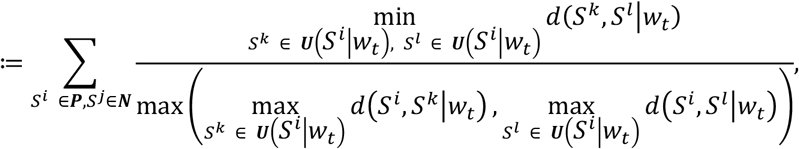

where ***P*** and ***N*** are the sets of sequences of known permissive and non-permissive orthologs, respectively, and ***U***(*S*^*i*^|*w*_*t*_) are the three orthologs closest to *S*^*i*^ in the distance matrix weighted by *w*_*t*_. Ideally, this score should be high when the distance matrix separates the pair of neighborhoods of most pairs of known permissive and non-permissive orthologs. The optimized weights are {*w*_*t*,1_ = 10, *w*_*t*,2_ = 10, *w*_*t*,3_ = 10}. Finally, we prioritized the orthologs using the following receptor score *R*(*S*^*i*^):

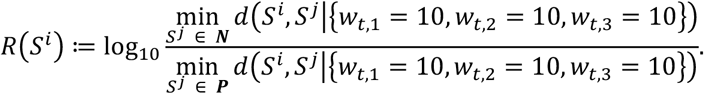

All orthologs with *R*(*S*^*i*^) ≥ 0.5 (candidate permissives) and *R*(*S*^*i*^) ≤ 0 (candidate non-permissives) were then prioritized (Table S2). All 11 candidates non-permissives except *Ornithorhynchus anatinus* and *Myotis davidii* (whose residue conformations at the S protein contact surfaces were extremely similar to that of human ACE2) were tested experimentally.

## Acknowledgements

This work was supported by the Beijing Nova Program of Science and Technology (https://mis.kw.beijing.gov.cn; no. Z191100001119064 to HC), the National Natural Science Foundation of China (http://www.nsfc.gov.cn; nos. 31801112, 61873276, 31900488 and 81830101 to HC, XB, HL and SW respectively), the Beijing Natural Science Foundation (http://kw.beijing.gov.cn/; no. 5204040 to HL), the Beijing Nova Program (https://mis.kw.beijing.gov.cn; no. Z171100001117119 to XW) and the Medical Innovation Project (no. 17SAZ13 to XW).

## Supplementary Data

**Supplementary Figure 1.**
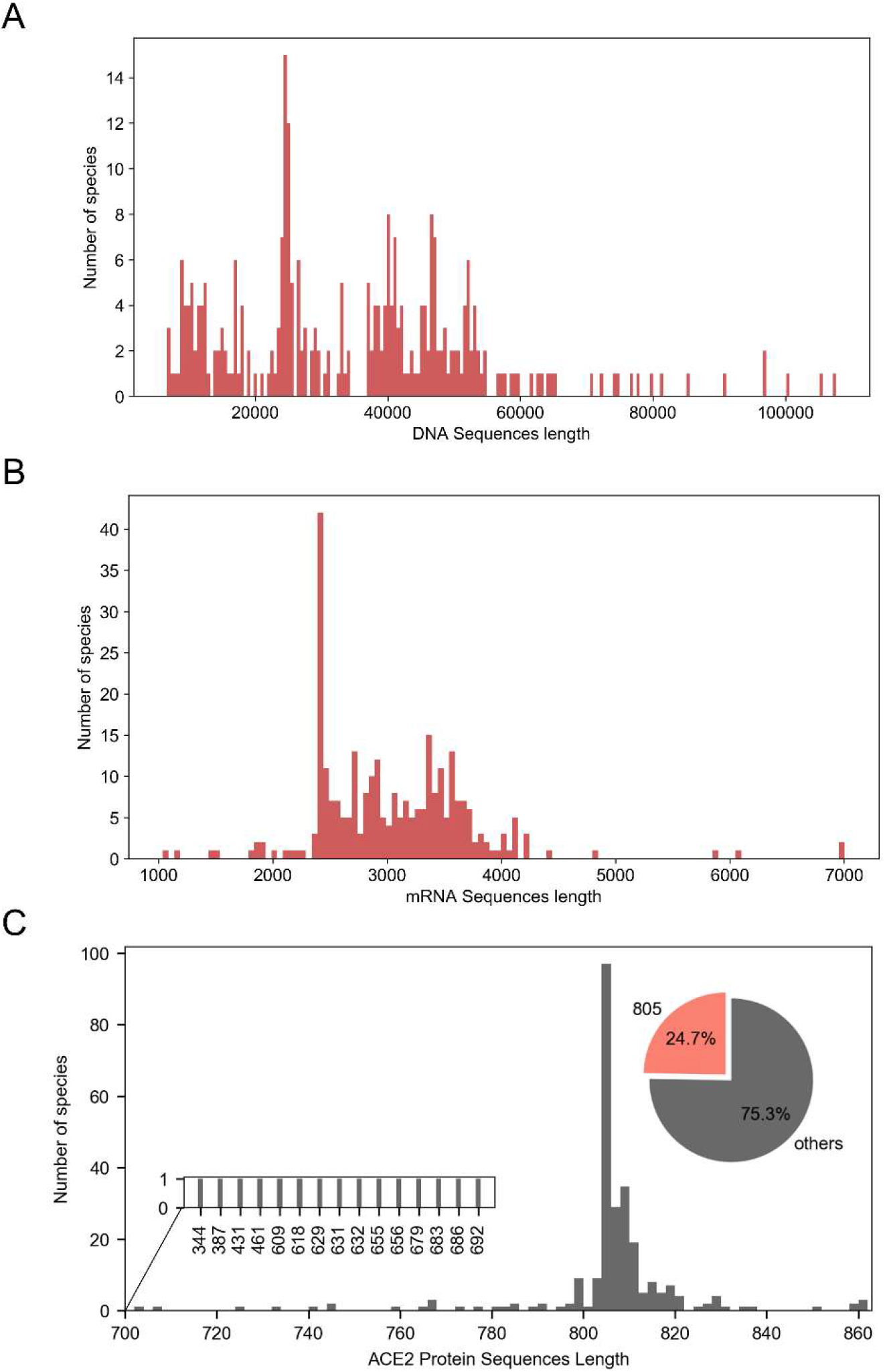
Frequency distributions of ACE2 DNA (A; bin width = 500) and mRNA (B; bin width = 50) sequence lengths in 286 vertebrates. (C) Frequency distribution of the ACE2 amino-acid sequence length in 287 vertebrates. Pie chart showing the relative proportions of species with the same ACE2 amino-acid lengths as humans and others. Bin width = 1.

**Supplementary Figure 2.**
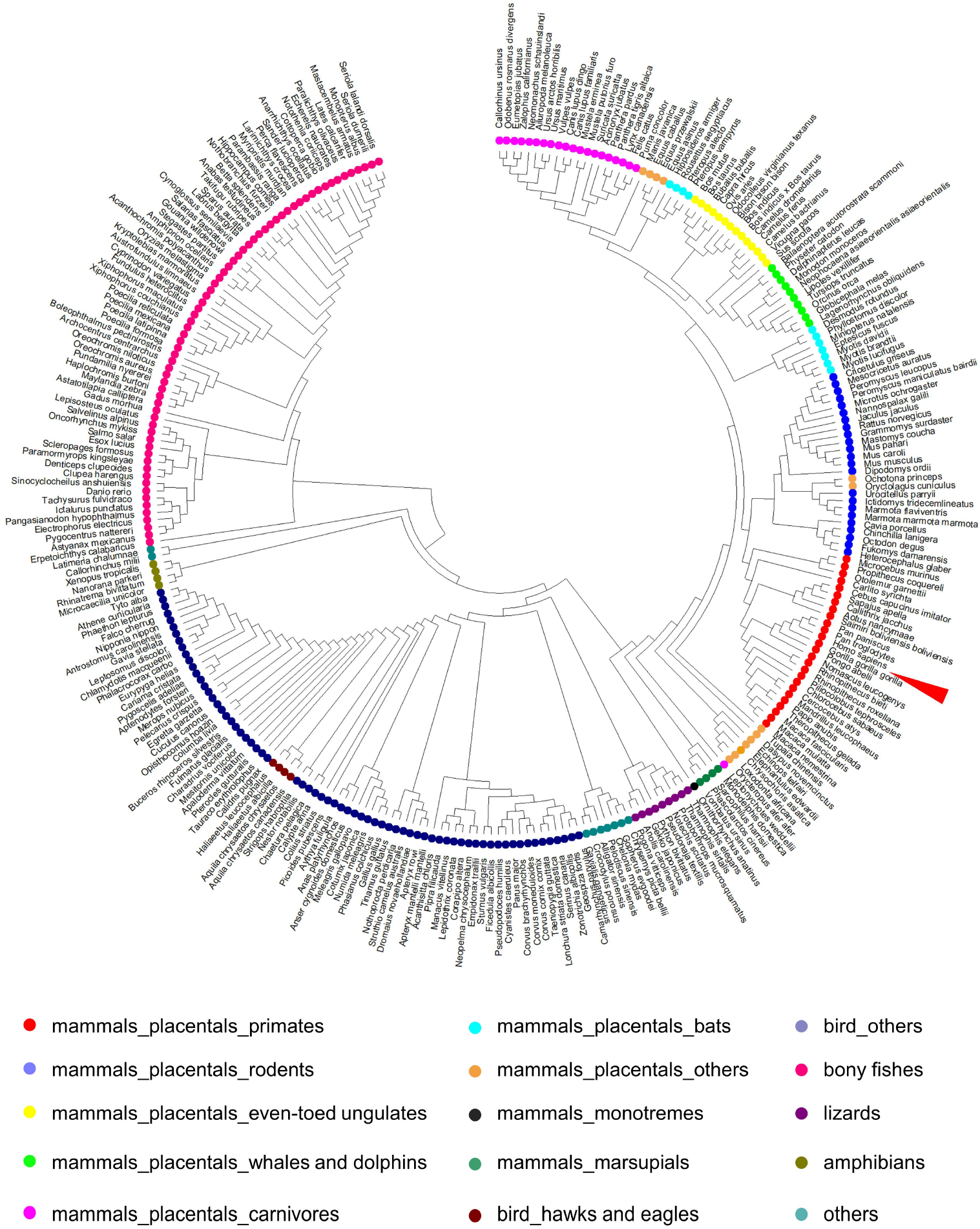
Evolution tree for whole ACE2 amino-acid sequences, built using the minimum-evolution method with the MEGA-X software (ver. 10.05) and the MUSCLE algorithm. Species of different taxa are represented by different colors.

**Supplementary Figure 3.**
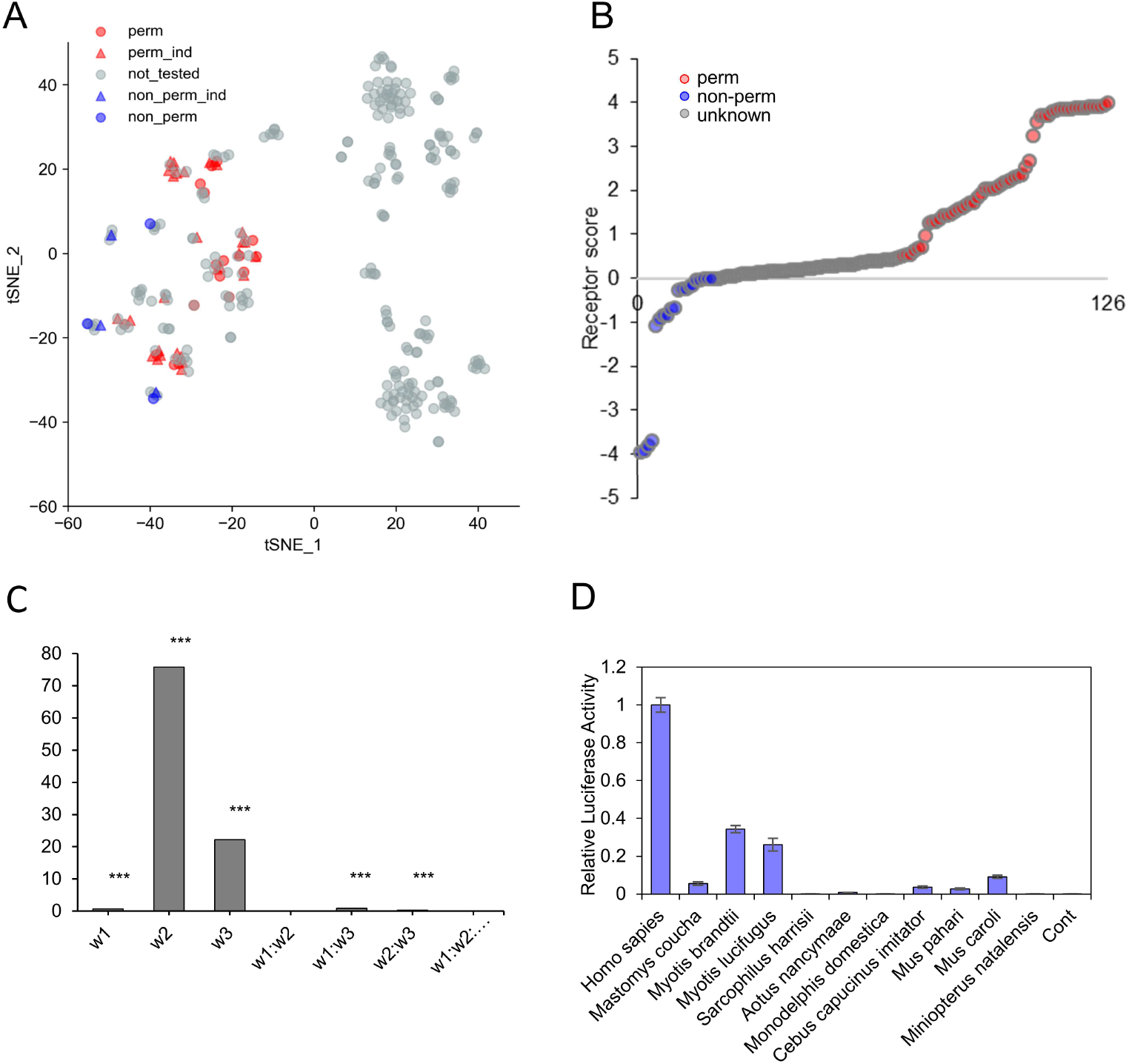
Evaluation of the residue-specific weighted calculation module. (A) t-SNE projection plot showing the clustering results for ACE2 orthologs from 287 vertebrates. perm, ACE2 orthologs permissive of SARS2pp entry (Figure 3); non_perm, ACE2 orthologs not permissive of SARS2pp entry (Figure 3); perm_ind, ACE2 orthologs permissive of SARS2pp entry, according to independent research^28^; non_perm_ind, ACE2 orthologs not permissive of SARS2pp entry, according to independent research^28^; not_tested, ACE2 orthologs that had never been tested. (B) Scatter plot of receptor scores for all mammals. (C) Top: schematic diagram of. Bottom: Multivariable variance results of residue-specific weighted calculation module. \*\*\**p* < 0.001. (D) Relative luciferase activity of different ACE2 orthologs in vector-transfected cells at 60 h after SARS2pp infection. Values are expressed as means with standard deviations (SDs; error bars).

**Supplementary Table 1.**
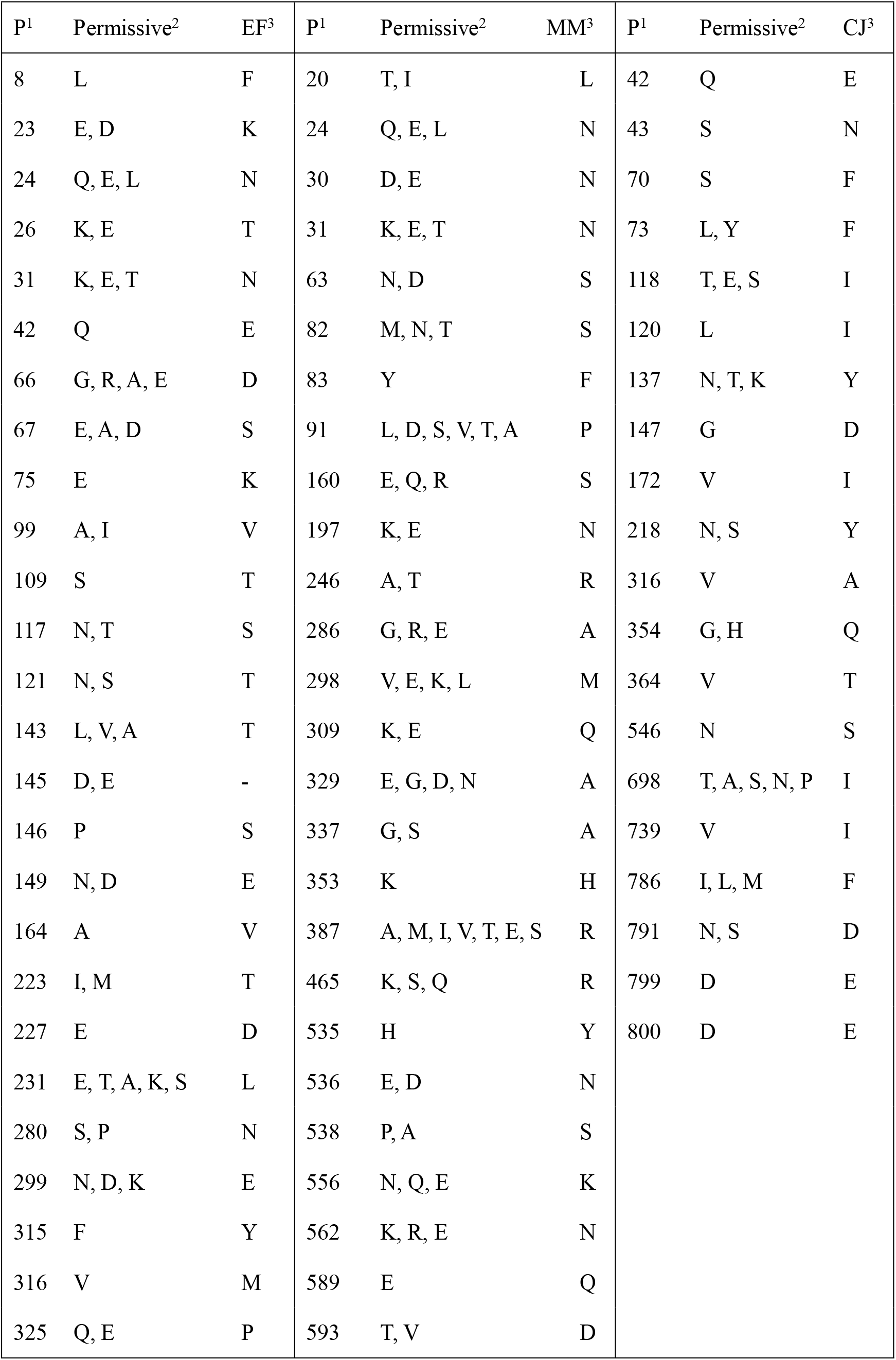
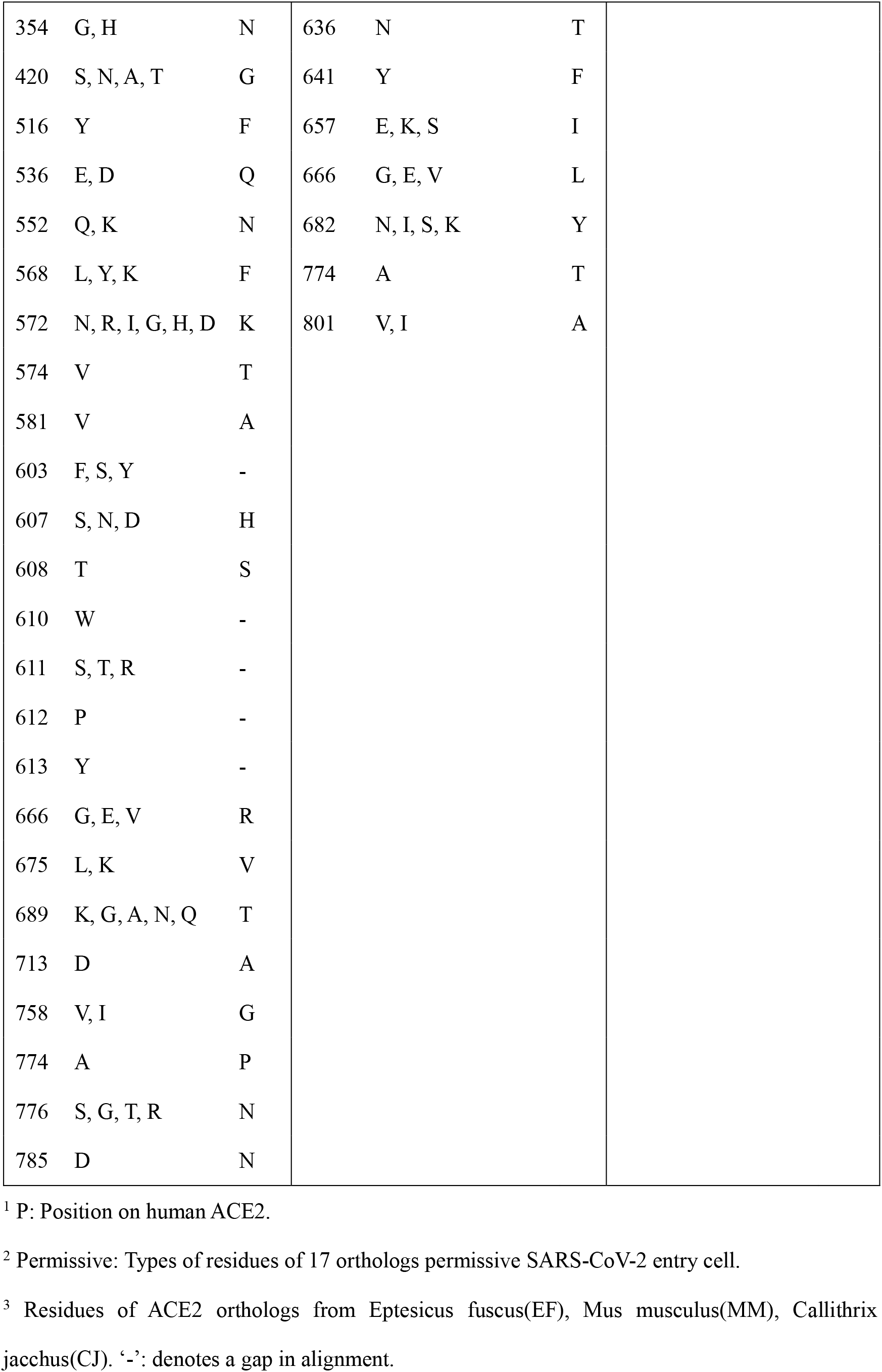
Key inconsistent residues between permissive and non-permissive orthologs

| P<sup>1</sup> | Permissive<sup>2</sup> | EF<sup>3</sup> | P<sup>1</sup> | Permissive<sup>2</sup> | MM<sup>3</sup> | P<sup>1</sup> | Permissive<sup>2</sup> | CJ<sup>3</sup> |
| --- | --- | --- | --- | --- | --- | --- | --- | --- |
| 8 | L | F | 20 | T, I | L | 42 | Q | E |
| 23 | E, D | K | 24 | Q, E, L | N | 43 | S | N |
| 24 | Q, E, L | N | 30 | D, E | N | 70 | S | F |
| 26 | K, E | T | 31 | K, E, T | N | 73 | L, Y | F |
| 31 | K, E, T | N | 63 | N, D | S | 118 | T, E, S | I |
| 42 | Q | E | 82 | M, N, T | S | 120 | L | I |
| 66 | G, R, A, E | D | 83 | Y | F | 137 | N, T, K | Y |
| 67 | E, A, D | S | 91 | L, D, S, V, T, A | P | 147 | G | D |
| 75 | E | K | 160 | E, Q, R | S | 172 | V | I |
| 99 | A, I | V | 197 | K, E | N | 218 | N, S | Y |
| 109 | S | T | 246 | A, T | R | 316 | V | A |
| 117 | N, T | S | 286 | G, R, E | A | 354 | G, H | Q |
| 121 | N, S | T | 298 | V, E, K, L | M | 364 | V | T |
| 143 | L, V, A | T | 309 | K, E | Q | 546 | N | S |
| 145 | D, E | - | 329 | E, G, D, N | A | 698 | T, A, S, N, P | I |
| 146 | P | S | 337 | G, S | A | 739 | V | I |
| 149 | N, D | E | 353 | K | H | 786 | I, L, M | F |
| 164 | A | V | 387 | A, M, I, V, T, E, S | R | 791 | N, S | D |
| 223 | I, M | T | 465 | K, S, Q | R | 799 | D | E |
| 227 | E | D | 535 | H | Y | 800 | D | E |
| 231 | E, T, A, K, S | L | 536 | E, D | N |  |  |  |
| 280 | S, P | N | 538 | P, A | S |  |  |  |
| 299 | N, D, K | E | 556 | N, Q, E | K |  |  |  |
| 315 | F | Y | 562 | K, R, E | N |  |  |  |
| 316 | V | M | 589 | E | Q |  |  |  |
| 325 | Q, E | P | 593 | T, V | D |  |  |  |

|  |  |  |  |  |  |
| --- | --- | --- | --- | --- | --- |
| 354 | G, H | N | 636 | N | T |
| 420 | S, N, A, T | G | 641 | Y | F |
| 516 | Y | F | 657 | E, K, S | I |
| 536 | E, D | Q | 666 | G, E, V | L |
| 552 | Q, K | N | 682 | N, I, S, K | Y |
| 568 | L, Y, K | F | 774 | A | T |
| 572 | N, R, I, G, H, D | K | 801 | V, I | A |
| 574 | V | T |  |  |  |
| 581 | V | A |  |  |  |
| 603 | F, S, Y | - |  |  |  |
| 607 | S, N, D | H |  |  |  |
| 608 | T | S |  |  |  |
| 610 | W | - |  |  |  |
| 611 | S, T, R | - |  |  |  |
| 612 | P | - |  |  |  |
| 613 | Y | - |  |  |  |
| 666 | G, E, V | R |  |  |  |
| 675 | L, K | V |  |  |  |
| 689 | K, G, A, N, Q | T |  |  |  |
| 713 | D | A |  |  |  |
| 758 | V, I | G |  |  |  |
| 774 | A | P |  |  |  |
| 776 | S, G, T, R | N |  |  |  |
| 785 | D | N |  |  |  |

<sup>1</sup> P: Position on human ACE2.
<sup>2</sup> Permissive: Types of residues of 17 orthologs permissive SARS-CoV-2 entry cell.
<sup>3</sup> Residues of ACE2 orthologs from *Eptesicus fuscus*(EF), *Mus musculus*(MM), *Callithrix* *jacchus*(CJ). ‘-’: denotes a gap in alignment.

**Supplementary Table 2.**
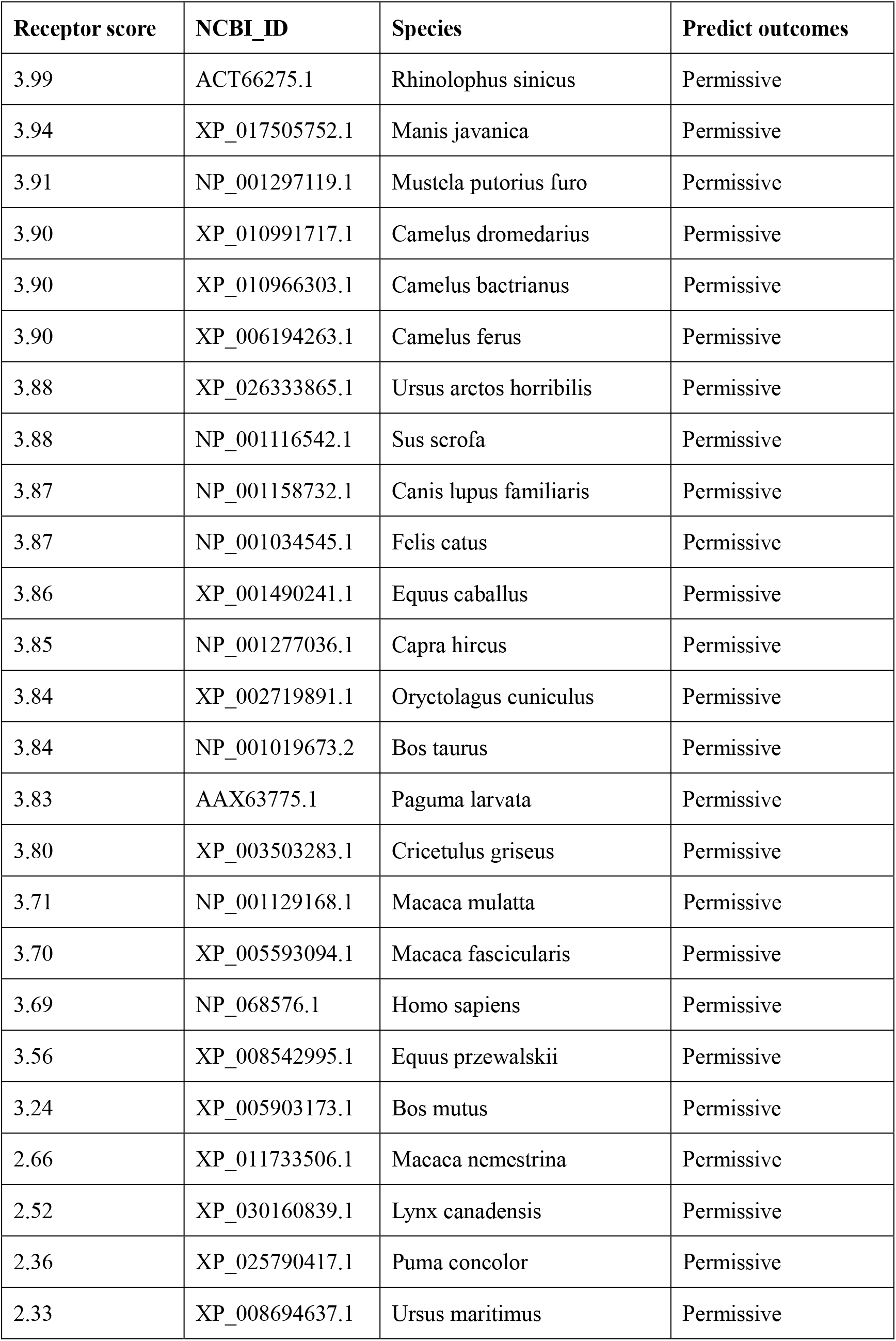
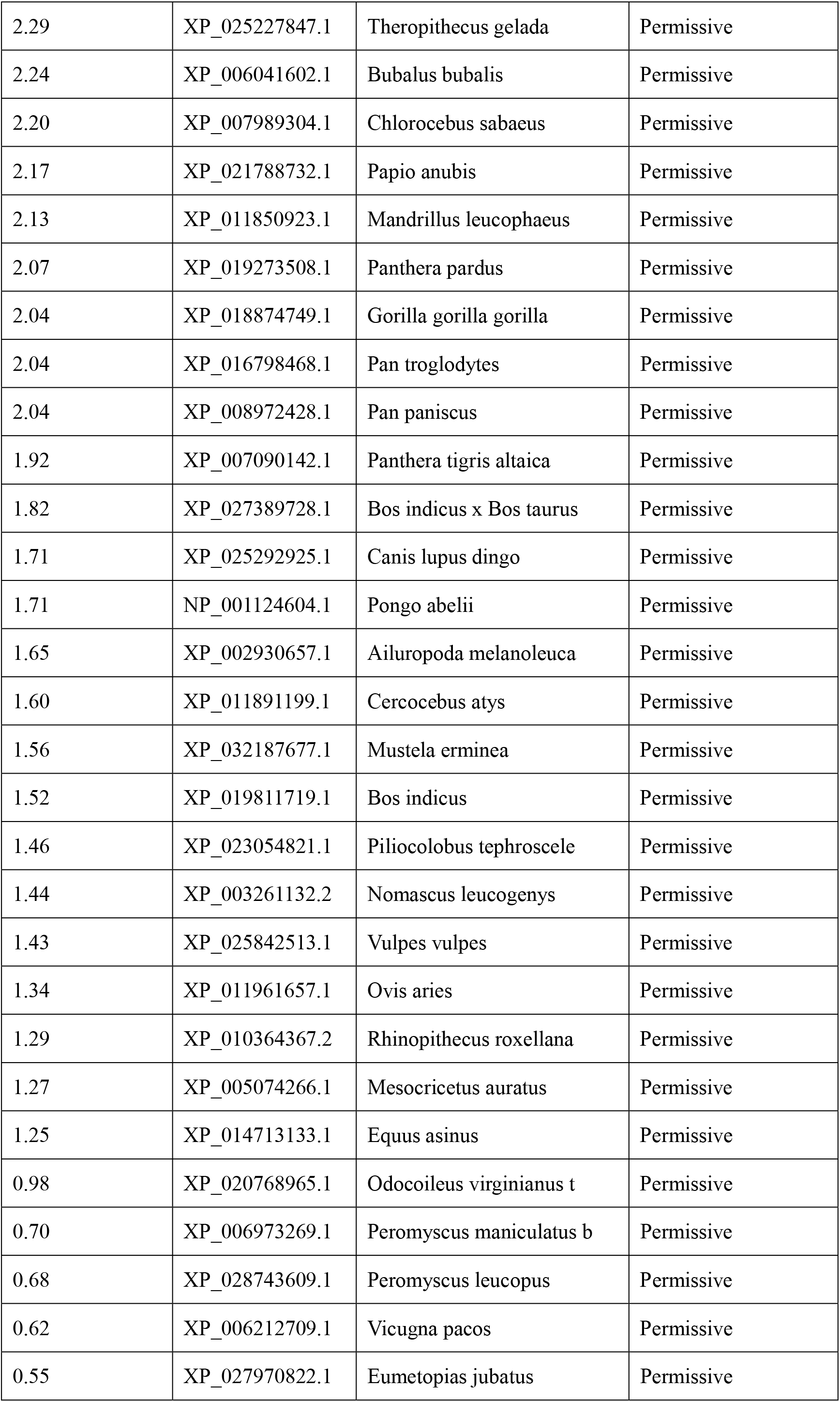
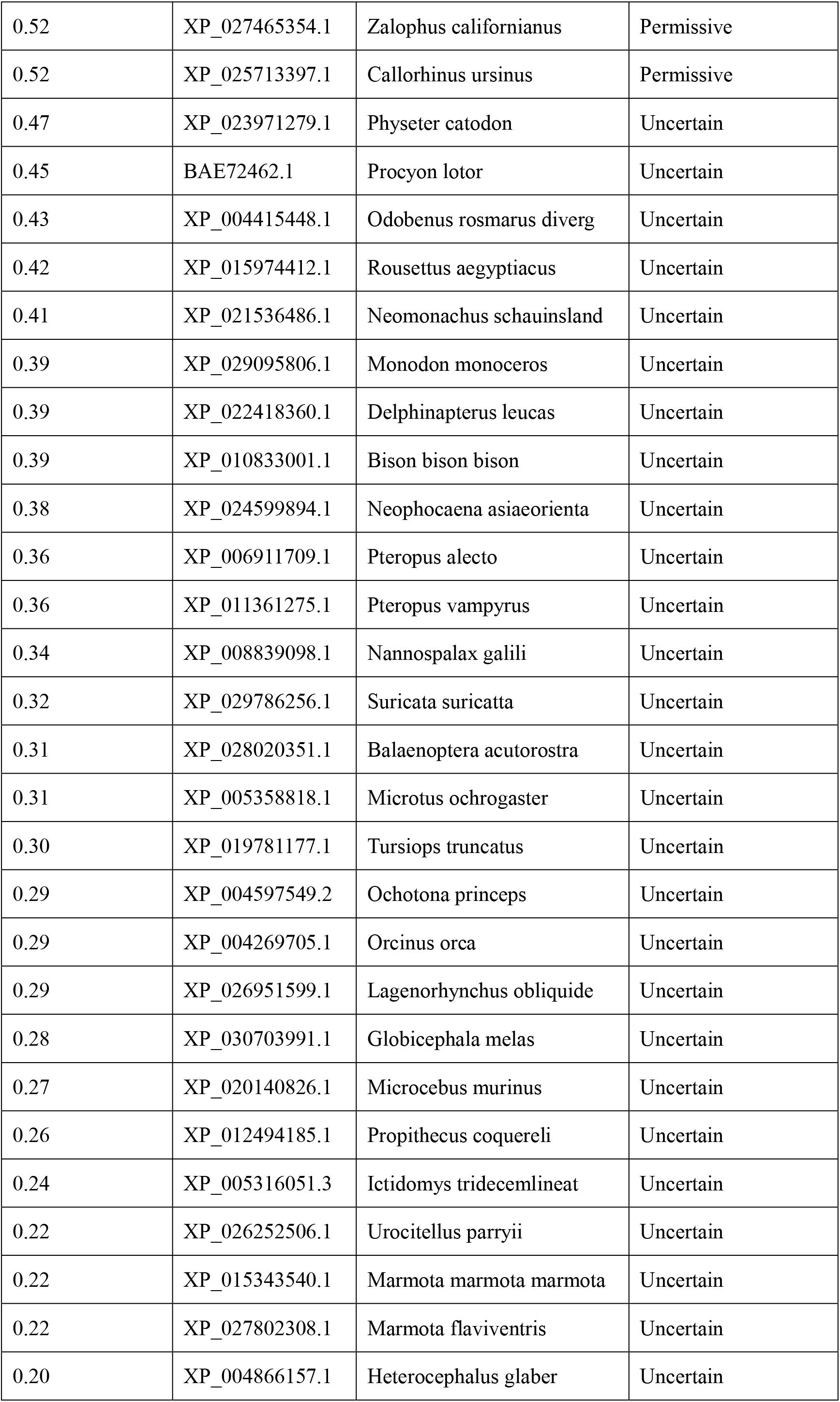
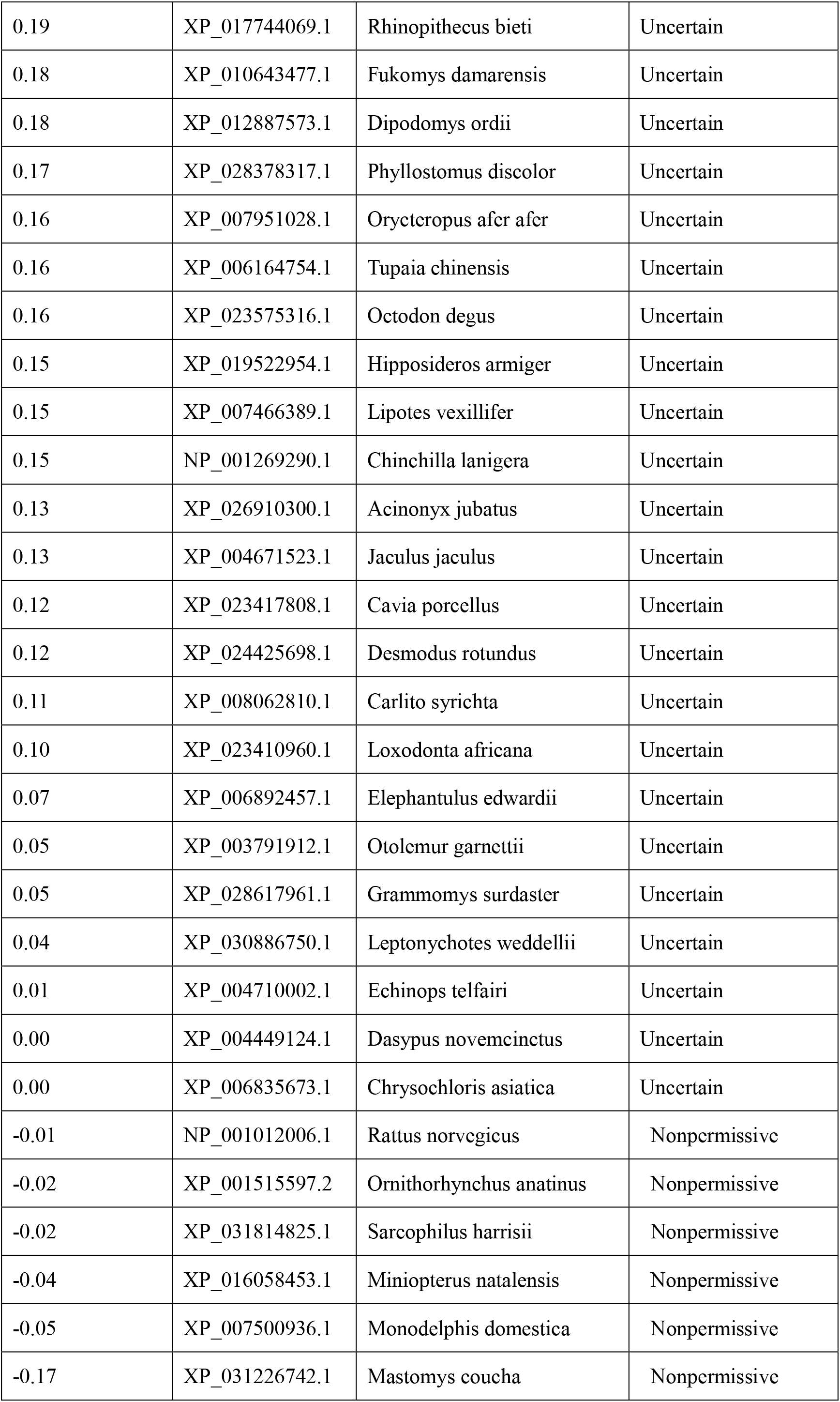
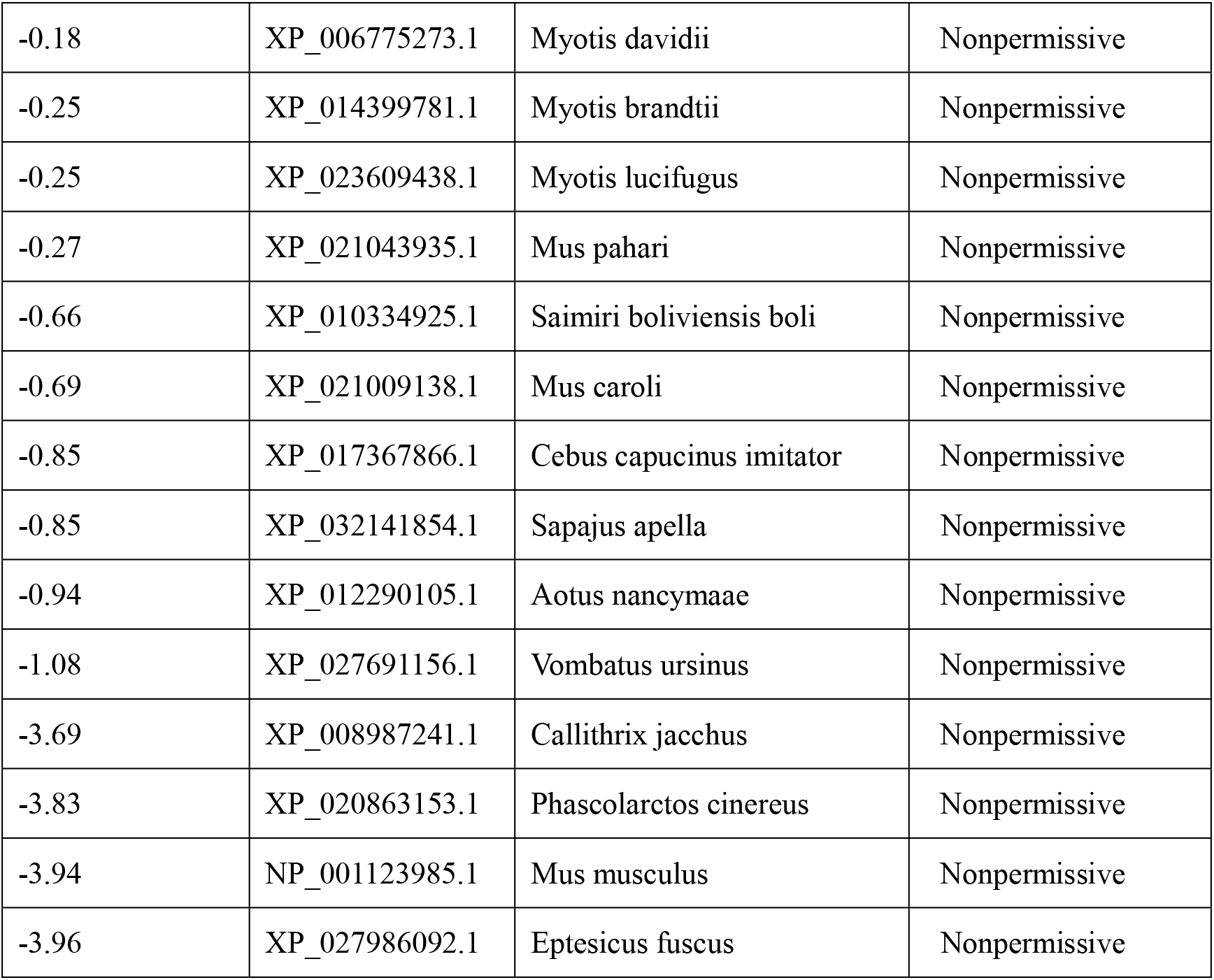
Predict outcomes of SARS-CoV-2 potential hosts

**Supplementary Table 3.** All 10×10×10=1,000 possible combinations of *w*_t_ ≔ {*w*_t,1_, *w*_t,2_, *w*_t,3_}

| Prior | Case | Possible weights |
| --- | --- | --- |
| $w_{t,1}$ | The $t$-th residue in the human ortholog is in the PD domain (19-615) | 1,2,3,4,5,<br>6,7,8,9,10 |
|  | Otherwise | 1 |
| $w_{t,2}$ | The $t$-th position is listed in Supplementary Table 1 (i.e., it is a key inconsistent residue between permissive and non-permissive orthologs ) | 1,2,3,4,5,<br>6,7,8,9,10 |
|  | Otherwise | 1 |
| $w_{t,3}$ | The $t$-th residue in the human ortholog is in the contact surface (PDB ID: 6M0J) of human ACE2 with S protein | 1,2,3,4,5,<br>6,7,8,9,10 |
|  | Otherwise | 1 |

